# Intrinsic persistent firing in CA1 encodes elapsed time across behaviorally relevant scales

**DOI:** 10.64898/2025.12.10.693586

**Authors:** Sara Zomorodi, Beate Knauer, Yacine Brahimi, Antonio Reboreda, Motoharu Yoshida, Zoran Tiganj

## Abstract

The ability to encode and maintain temporal relationships is crucial for learning, predicting, and forming episodic memories. While hippocampal time cells and entorhinal temporal context cells are well-established *in vivo*, it remains unclear whether single neurons can sustain representations of elapsed time over multi-second intervals, independent of synaptic drive. Here, using whole-cell patchclamp recordings in rat hippocampal CA1 slices under synaptic blockade, we show that following a brief current pulse, many neurons exhibit exponentially decaying firing rates with a broad distribution of time constants extending to tens of seconds. This indicates that single neurons possess intrinsic mechanisms capable of covering behaviorally relevant temporal scales. These results highlight that single neurons can encode temporal information over more than an order of magnitude, providing a potential cellular substrate for temporal coding in the brain.

## 1 Introduction

Our ability to remember what happened and when is critical for learning temporal relationships and predicting future outcomes. A wealth of evidence implicates the hippocampal formation as a central hub for encoding and retrieving temporal information (e.g., Fortin et al. (2002); Lehn et al. (2009); Howard and Eichenbaum (2013); Hsieh et al. (2014); Eichenbaum (2014)).

Seminal *in vivo* studies identified hippocampal “time cells”, neurons that fire sequentially after salient events, providing a timeline for temporally organized memories (Eichenbaum, 2014; MacDonald et al., 2011; Pastalkova et al., 2008; Umbach et al., 2020). Complementing this sequential activity, neurons named “temporal context cells” observed in the lateral entorhinal cortex (LEC), a region projecting to the hippocampus, were found to exhibit gradually changing firing patterns (Tsao et al., 2018; Bright et al., 2020). Such ramping cells were also reported in other regions, including the prefrontal cortex and posterior parietal cortex (Kim et al., 2013; Morcos and Harvey, 2016; Narayanan, 2016; Cao et al., 2024; Affan et al., 2025).

While both sequential time cells and gradually changing temporal context cells are well-established in the behaving animal, the underlying mechanisms that enable neurons to represent time far beyond the duration of a stimulus remain unresolved. In particular, it is unclear whether these temporal dynamics necessitate only network-level interactions or if intrinsic properties of individual neurons are also involved.

Computational work suggests that network mechanisms based on recurrent activity (e.g., reservoir or attractor dynamics) can give rise to both temporal context cells and sequential time cells (Zylberberg and Strowbridge, 2017; Masse et al., 2019; Voelker and Eliasmith, 2018). Modeling work also suggests that single cells could intrinsically generate exponentially decaying firing rates to encode elapsed time (Tiganj et al., 2015; Liu et al., 2019). A population of such neurons with a spectrum of decay rates could be linearly transformed to produce the sequential activity characteristic of time cells, akin to an inverse Laplace transform (Shankar and Howard, 2012; Howard et al., 2014; Tano et al., 2020).

The biological plausibility of an intrinsic cellular mechanism is supported by *in vitro* studies demonstrating that, in the presence of synaptic blockers, individual neurons in the hippocampus and entorhinal cortex can generate persistent firing following a brief stimulus, often induced by cholinergic modulation (Klink and Alonso, 1997; Fransén et al., 2006; Yoshida and Alonso, 2007). However, it is not well understood whether this intrinsically sustained activity can carry graded temporal information. Specifically, it remains unknown whether single neurons can produce gradually decaying firing rates across the wide range of timescales relevant for temporal coding observed *in vivo*. Such long timescales could play an important role in bridging events in behavioral tasks.

To address this gap, we investigate whether individual hippocampal CA1 neurons, isolated from their network, can intrinsically encode temporal information. We used in *vitro* whole-cell recordings in the presence of synaptic blockers and the cholinergic agonist carbachol (CCh). Neurons were stimulated with brief current injections to induce persistent firing. We demonstrate that many individual CA1 neurons exhibit a gradually decaying firing rate that is well-described by an exponential function. Critically, the time constants of this decay span a wide, continuous range, from seconds to tens of seconds. These results suggest that single neurons possess intrinsic mechanisms to encode temporal information over behaviorally relevant scales, offering a fundamental cellular substrate for memory-encoding circuits.

## 2 Methods

### 2.1 Animals

The experiments were performed at the Ruhr University Bochum, Bochum, Germany, between 2011 and 2014. Experiments were conducted following the guidelines of the local animal ethics committees and the European Communities Council Directive of September 22, 2010 (2010/63/EU). All experimental protocols were approved by the local ethics committee (Der Tierschutzbeauftragte, Ruhr-Universität Bochum). 14-24 days old male and female Long-Evans rats were used to collect data from hippocampal CA1, using three different concentrations of CCh (5,10, 20 *μ*M). The protocols have been previously described in detail (Knauer and Yoshida, 2019).

### 2.2 Acute brain slice preparation

The rats were deeply anesthetized with an intraperitoneal injection of a ketamine:xylazine (100:4 mg/kg) cocktail. Transcardial perfusions with ice-cold cutting solution containing (in mM) 110 choline chloride, 1.25 NaH_2_PO_4_, 7 MgCl_2_, 2.5 KCl, 7 D-Glucose, 3 pyruvic acid, 1 ascorbic acid, 26 NaHCO_3_, and 0.5 CaCl_2_ were performed. The brains were quickly removed from the cranial cavities and immersed in ice-cold cutting solution. 350 *µ*m horizontal slices were obtained using a vibratome (VT1000 S, Leica Instruments). Brain slices were transferred into a holding chamber filled with artificial cerebrospinal fluid (ACSF) containing (in mM) 125 NaCl, 1.2 NaH_2_PO_4_, 1.8 MgSO_4_, 3 KCl, 10 D-glucose, 26 NaHCO_3_, and 1.6 CaCl_2_. The slices were subsequently held for 30 min at 30 °C. The pH of the cutting solution and the ACSF were constantly maintained at 7.4 by saturation with 95% O_2_–5% CO_2_. Slices were kept at room temperature for at least 30 min before recording.

### 2.3 Recording

The slices were then transferred to a recording chamber perfused with oxygenated ACSF (35 ± 1 °C) supplemented with kynurenic acid (2 *mM*) and picrotoxin (0.1 *mM*) to block fast ionotropic glutamate and GABA_A_ synaptic transmission, respectively. The cells were visualized using an upright microscope (Axioscope, Zeiss) equipped with a 4x objective, a 40x water-immersion objective, and a monochrome camera (WAT-902H Ultimate, Watec) connected to a computer. The CA1 region was targeted using the 4x objective lens, and the cells were identified with the 40x lens. Patch pipettes (3–8 MΩ) were pulled from borosilicate glass capillaries (GB150-8P, Science Products) using a P-87 horizontal puller (Sutter Instruments). The pipettes were back-filled with filtered intracellular solution containing (in mM) 120 K-gluconate, 10 HEPES, 0.2 EGTA, 20 KCl, 2 MgCl_2_, 7 PhCreat di(tris), 4 Na_2_ATP, 0.3 Tris-GTP and 0.1% biocytin for post-hoc location verification (pH adjusted to 7.3 with KOH). The whole-cell patch configuration was achieved by forming a tight seal (≥ 1 GQ) on the somata of the cells, and electrical access was achieved by applying negative pressure. Electrical signals were amplified using an AxoClamp-2A amplifier (Axon Instruments) and a Multiclamp 700 B (Axon Instruments). The signals were sampled at 20 kHz and low-pass filtered at 10 kHz. Recordings were obtained in current-clamp mode using Clampex 9.0 data acquisition software. The liquid junction potential was not corrected.

### 2.4 Induction of persistent firing

Prior to inducing persistent firing, the membrane potential was adjusted to slightly below the level where spontaneous spikes occur using DC current injection while the cholinergic receptor agonist carbachol (5, 10 or 20 *μ*M) was continuously bath applied (Knauer et al., 2013). Persistent firing was then induced using a brief current injection (2 s, 100 pA). The membrane potential recorded after the offset of the brief current injection was used to analyze persistent firing in the following sections.

### 2.5 Data analysis

We first transformed the raw membrane-potential recordings into instantaneous firing rates. An action potential was identified when the membrane potential surpassed a threshold of –10 *mV*, indicating a depolarization event. Binned firing rates were computed as the number of spikes occurring within 1 *s* windows, advanced in 100 ms steps.

To characterize gradual changes in neuronal activity following the current injection, we fitted exponential and linear functions to the binned firing rate. Specifically, to test the hypothesis about exponentially changing firing rate, we modeled the firing rate of individual neurons using the following model:

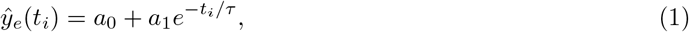

with parameters *a*_*0*_ and *a*_*1*_ capturing the magnitude of the firing rate, and *τ* capturing the time constant. To keep *ŷ* strictly positive, we constrained the parameters such that *a*_*0*_ *>* 0 and *a*_*0*_ + *a*_*1*_ *>* 0.

To evaluate whether linearly changing firing rate could better describe the neuronal activity, we fitted the firing rate with the following model:

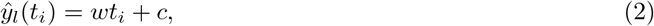

that included slope parameter *w* and intercept parameter c. We also computed a fit with a constant term *ŷ*_*c*_*(t*_*i*_) = c, which captures neurons that responded to the current injection with stable persistent firing.

We fit 427 bins (≈ 43.5 s) beginning at the offset of the current pulse. The parameters were optimized to reduce root mean square error (RMSE) between *ŷ* and the true firing rate y(t):

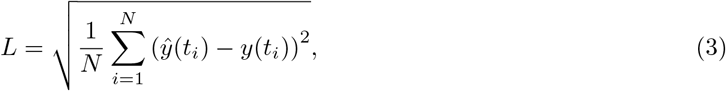

where *N* = 427 is the number of time bins, *t*_1_ =0 *s* and *t*_427_ ≈ 43 *s*. Curve-fitting was done using the Python package *lmfit*.

For each cell, we compared adjusted R^2^ between the exponential, linear and constant models:

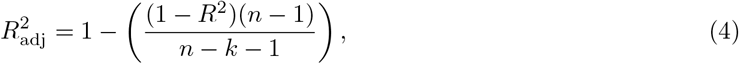

where *n* is the number of time bins and *k* is the number of free parameters (*k* = 3 for exponential model, *k* = 2 for linear model and *k* = 1 for the constant model) and R^2^ is coefficient of determination:

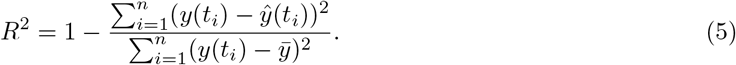

To evaluate the robustness of the temporal changes in the persistent firing of exponentially decaying cells, we estimated the elapsed time since the current injection from each neuron:

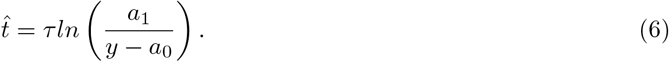

Because our central question was whether monotonic temporal kernels can arise intrinsically, model comparison was restricted *a priori* to exponential, linear, and constant functions. We therefore do not interpret the excluded neurons as exhaustively classified. Other motifs, including biphasic, delayed-peak, oscillatory, or burst-like activity under cholinergic drive, may also occur in this preparation.

For anatomical analyses, soma position was reconstructed from biocytin-filled neurons in horizontal slices. Proximo-distal position was expressed as a percentage distance within CA1, with 0% defined at the CA1/CA2 border and 100% defined at distal CA1 toward the subiculum. Relative longitudinal (dorsal-ventral) position was estimated using the same slice-based anatomical procedure described in Knauer and Yoshida (2019): the serial number of the slice during cutting, whether the recording was obtained from the top or bottom side of the slice, and the measured amount of remaining brain tissue after slicing were used to estimate the cell’s position in millimeters from the top of the brain. The zero point corresponds to approximately 2 mm below bregma and a larger dorso-ventral distance corresponds to a more ventral position. Because the slices sampled only a limited portion of the longitudinal axis, this measure should be interpreted as a relative position within the sampled horizontal slice series rather than as a full septotemporal coordinate across the entire hippocampus.

## 3 Results

We analyzed *in vitro* activity from 98 CA1 pyramidal neurons across three different CCh concentrations following a current pulse injection. We focused on neurons with sustained gradually changing firing rates, which did not self-terminate for at least 30 s following the pulse injection (Fig. 1A-F).

**Figure 1:**
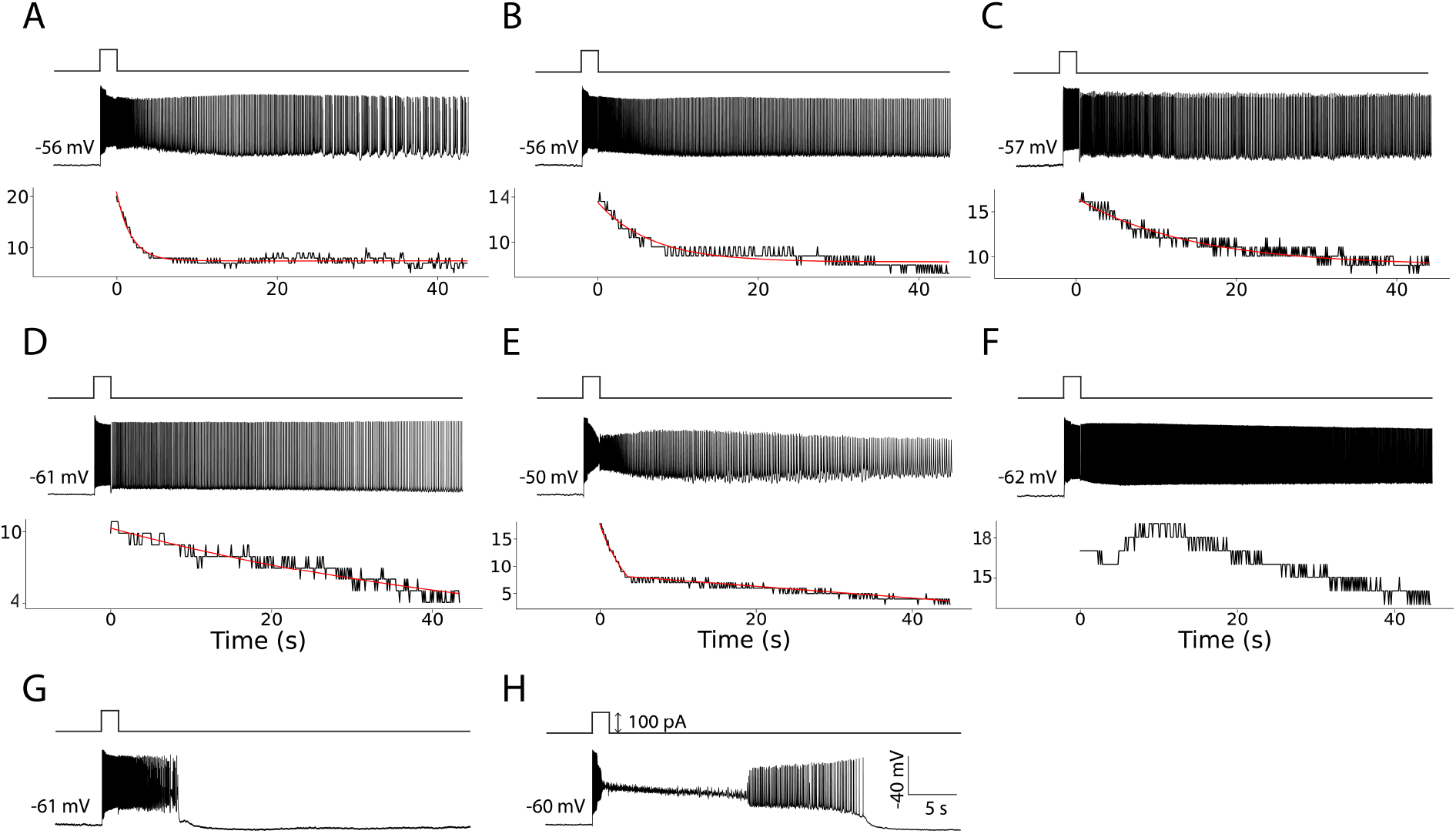
Examples of neurons with: **A.-C.** Exponentially decaying firing rate, sorted by time constant from smallest (left) to largest (right). **D.** A firing rate best fit with a linear function. **E.** A firing rate best fit with a combination of linear and exponential fits. **F.** Maximum firing rate occurring after 5 *s*. **G.** Self-terminating persistent firing **H.** Depolarization block. In each panel, the top row shows the time when the current was injected, the second row shows raw membrane potentials, and the third row (if applicable) shows firing rate (Hz) (black lines) and the best fit with either an exponential or a linear function (red lines).

We considered that depolarization block had occurred when the membrane potential entered a sustained depolarized plateau above the cell’s physiological spike threshold, accompanied by a loss of overshooting spikes (i.e., no events crossing –10 *mV*) for more than 5 s after current injection. These criteria for self-termination and depolarization block were adopted from Brahimi et al. (2023). We did not observe any neurons for which the fit with a constant term was better than the fit with either an exponential or a linear model. Thus, out of 98 neurons, the firing activity of 67 (68%) neurons was best fit with models that include gradual change across time.

### 3.1 Exponentially decaying firing was the most commonly observed activity profile

Among the 67 remaining neurons with gradually changing firing activity, 32 neurons (48%) were best fit with the exponential decay (Fig. 1A-C). We note that in all cases where the exponential model was better than others, the exponent was always negative, indicating a decaying firing rate. We also observed 11 neurons (16%) where the linear fit was better than the exponential one (Fig. 1D, Table 1), also with a negative slope indicating again a decaying firing rate.

**Table 1:** Response types of CA1 pyramidal neurons at three CCh concentrations. Numbers indicate counts of neurons in each category. **DB**, depolarization block; **ST**, self-terminating within 30 s; **Peak**, delayed peak (> 5s after pulse); **Two-phase**, biphasic decay with a rapid initial drop followed by a slower phase. **Lin. dec**., linearly decaying firing; **Exp. dec**., exponentially decaying firing;

| CCh (μM) | # Total | # DB | # ST | # Peak | # Two-phase | # Lin. dec. | # Exp. dec. |
| --- | --- | --- | --- | --- | --- | --- | --- |
| 5 | 25 | 1 | 3 | 2 | 3 | 3 | 13 |
| 10 | 39 | 2 | 4 | 6 | 7 | 5 | 15 |
| 20 | 34 | 13 | 8 | 4 | 2 | 3 | 4 |

A subset of 24 neurons was poorly described by both exponential and linear models and was therefore excluded from these counts and the subsequent analysis. Specifically, some neurons whose activity was nominally best captured by the exponential model exhibited two phases: a steep decline during the first ∼ 5 *s* followed by a slower decay. While the exponential fit was better than the linear or constant fit for such cells, it was overall poorly capturing the firing dynamics (Fig. 1E). To identify these cells, we separated the neuronal activity into two sections, before and after the first 5 *s*, and compared the adjusted R^2^ for a combination of linear and exponential fits for each section. This new fit had a higher adjusted R^2^ (with 5 parameters) than the exponential fit for 12 neurons (17%). In addition, 12 neurons were fitted best with decaying functions, but had a temporary increase in firing activity that peaked more than 5 s after the current injection (Fig. 1F). We consider the firing rate of these neurons as non-monotonically changing rather than gradually increasing or decreasing. Activity of all analyzed neurons together with exponential and linear fits is shown in the Supplementary Material.

Of the 98 neurons, 31 of them did not show such persistent firing activity due to either self-terminating activity (Fig. 1G, 15 neurons) or depolarization block (Fig. 1H, 16 neurons) (Table 1). Since neurons whose activity was best fit with the exponentially decaying functions were by far most frequent, they were the focus of our subsequent analysis, where we further investigated their firing properties.

### 3.2 Diversity of time scales: exponential decay spans a range of time constants extending to tens of seconds

We observed a broad range of temporal scales for neurons whose activity was best fit with the exponentially decaying model. The distribution of fitted time constants is shown in Fig. 2A. Fig. 2B-D show the histograms for each of the three CCh concentrations. The time constants overall ranged from 1.1 *s* to 49.3 *s*. To provide a better sense of the activity profiles, plots in the Supplementary Material show activity of all 32 neurons together with the exponential fits sorted by the time constant of the fit (for illustrations see Fig. 1A-C for neurons fitted with exponentially decaying functions with time constants of 1.81 *s*, 6.61 *s* and 14.12 *s* respectively). In general, the number density of neurons decreased as a function of time constant, contributing to the gradual decrease of temporal resolution following the current injection. We found no effect of carbachol concentration on the decay time constants. A Kruskal-Wallis test showed no significant difference among the three groups (5, 10, and 20 *µ*M; *H*(2, *N*=32) = 0.41, *p* = 0.81, *η*^2^ = 0.02). However, we note that the proportion of neurons exhibiting such decay was significantly lower at 20 *µM* (Table 1), suggesting that while higher cholinergic drive reduces the prevalence of this coding regime, it may not alter the temporal dynamics of the cells that remain within it.

**Figure 2:**
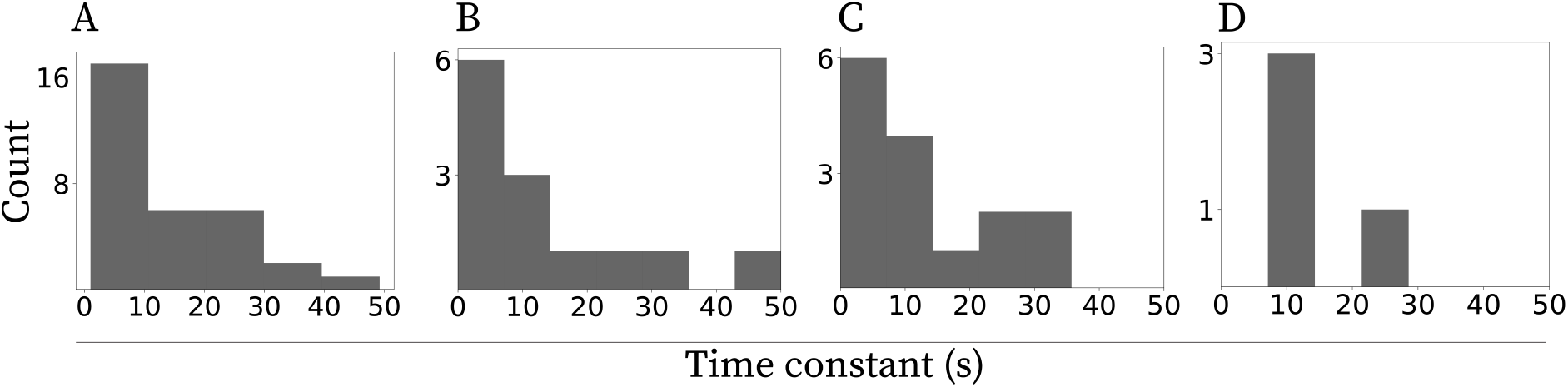
Neurons with exponentially decaying firing profiles span a wide range of temporal scales. Histograms of time constants indicate a decreasing number density of neurons as a function of the time constant. **A.** Neurons from all three CCh concentrations combined. **B.** 5 *µ*M CCh. **C.** 10 *µ*M CCh. **D.** 20 *µ*M CCh.

### 3.3 Higher CCh concentration shifts the activity profiles away from exponential decay and toward DB and/or ST

We compared the proportion of CA1 neurons whose firing was best fit with the exponentially decaying model across three carbachol concentrations (5, 10, and 20 *µ*M; 25, 39, and 34 neurons, respectively). A 𝒳^2^ test of independence revealed a significant association between concentration and decay type, *𝒳*^*2*^(2,N=98) = 11.60, p = 0.0030, Cramér’s *V* = 0.34 (moderate effect).

Pairwise Fisher exact tests (Holm-corrected) showed that the 20 *µM* group had significantly fewer exponentially decaying neurons than both 5 *µM* (odds ratio = 8.1, p_corr_ = 0.0035) and 10 *µM* (odds ratio = 4.7, p_corr_ = 0.030), whereas the difference between 5 *μ*M and 10 *μ*M was not significant (p_corr_ = 0.31). Overall, our results indicate that exponentially decaying firing is more prevalent at lower CCh concentrations. Beyond the reduction in exponentially decaying neurons, higher CCh also shifted response outcomes toward failure modes. At 20 *μ*M, the proportion of depolarization block (DB) rose to 38.2% (13/34) compared with 4.0% (1/25) at 5 *μ*M and 5.1% (2/39) at 10 *μ*M; self-terminating (ST) responses likewise increased to 23.5% (8/34) vs. 12.0% (3/25) and 10.3% (4/39). Fisher’s exact tests comparing 20 *μ*M to pooled 5–10 *μ*M indicated substantially higher odds of DB (odds ratio OR = 12.59, *p* = 4.39 × 10^−5^) and a moderate, non-significant increase for ST (OR = 2.51, p = 0.140). These patterns suggest that excessive cholinergic drive increases the likelihood that persistent firing either collapses (ST) or enters a depolarized, spike-incompetent state (DB).

### 3.4 Exponential decay can enable estimation of elapsed time

Figure 3 shows estimation of elapsed time from the exponentially decaying neurons. The end of the current injection is considered time zero. The estimation was done for each neuron with exponentially decaying firing rate following Eq. 6. We then averaged the estimated time from all neurons from the same CCh condition and compared it to the actual elapsed time. Consistent with Weber’s law, the standard deviation of the estimate increased with elapsed time. The correlation between standard deviation of the estimated time and the real time was found to be significant for all conditions (for 5 *µ*M CCh *r* =0.849,*p* < 0.001, for 10 *µ*M CCh *r* =0.909,*p* < 0.001, for 20 *µ*M CCh *r* =0.556,*p* < 0.001). Note that since this analysis was conducted with a different number of neurons per condition, the statistical power in each condition was rather different (Table 1), accounting for the large variability observed across conditions.

**Figure 3:**
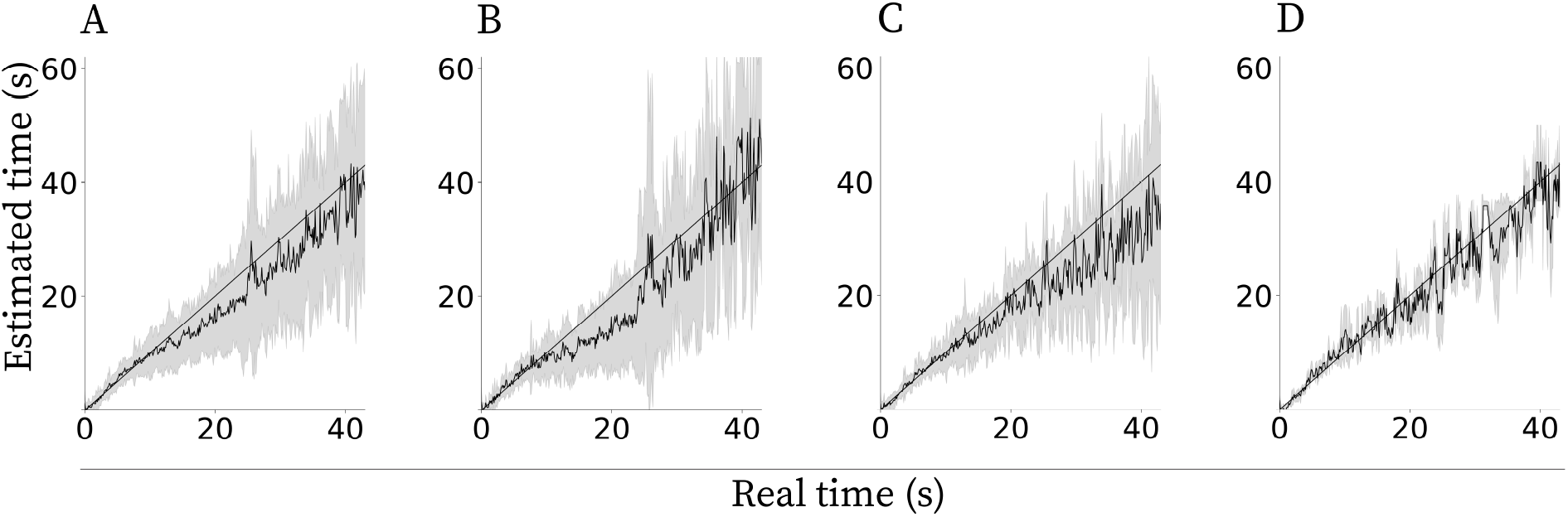
Estimating elapsed time from exponentially decaying cells. Note that the standard deviation of the estimate (gray lines) increases with time. A solid straight line represents a perfect estimate and a solid curvy line is the mean estimate across cells. **A**. Neurons from all three CCh concentrations combined. **B**. 5 *μ*M CCh. **C**. 10 *μ*M CCh. **D**. 20 *μ*M CCh.

### 3.5 Anatomical position and age were not detectably related to decay time constants

To test whether decay time constants vary systematically with cell location and age, we pooled neurons across all CCh concentrations and correlated each neuron’s time constant with its relative longitudinal (dorsal–ventral) position within the sampled horizontal slice series and its proximo–distal position within CA1 (Figure 4), as well as animals’ age (14–24 days). After Bonferroni correction across three comparisons, Pearson coefficients were not significant. Specifically for dorsal-ventral: r = −0.28, p = 0.30, *n* = 32, for proximo–distal: *r* = −0.40, *p* = 0.06, *n* = 32 and age *r =* 0.36, *p =* 0.04, *n =* 32. Together, these analyses indicate that neither recording depth, proximo–distal position, nor age measurably influences the range of decay time constants observed in CA1 neurons under our conditions.

**Figure 4:**
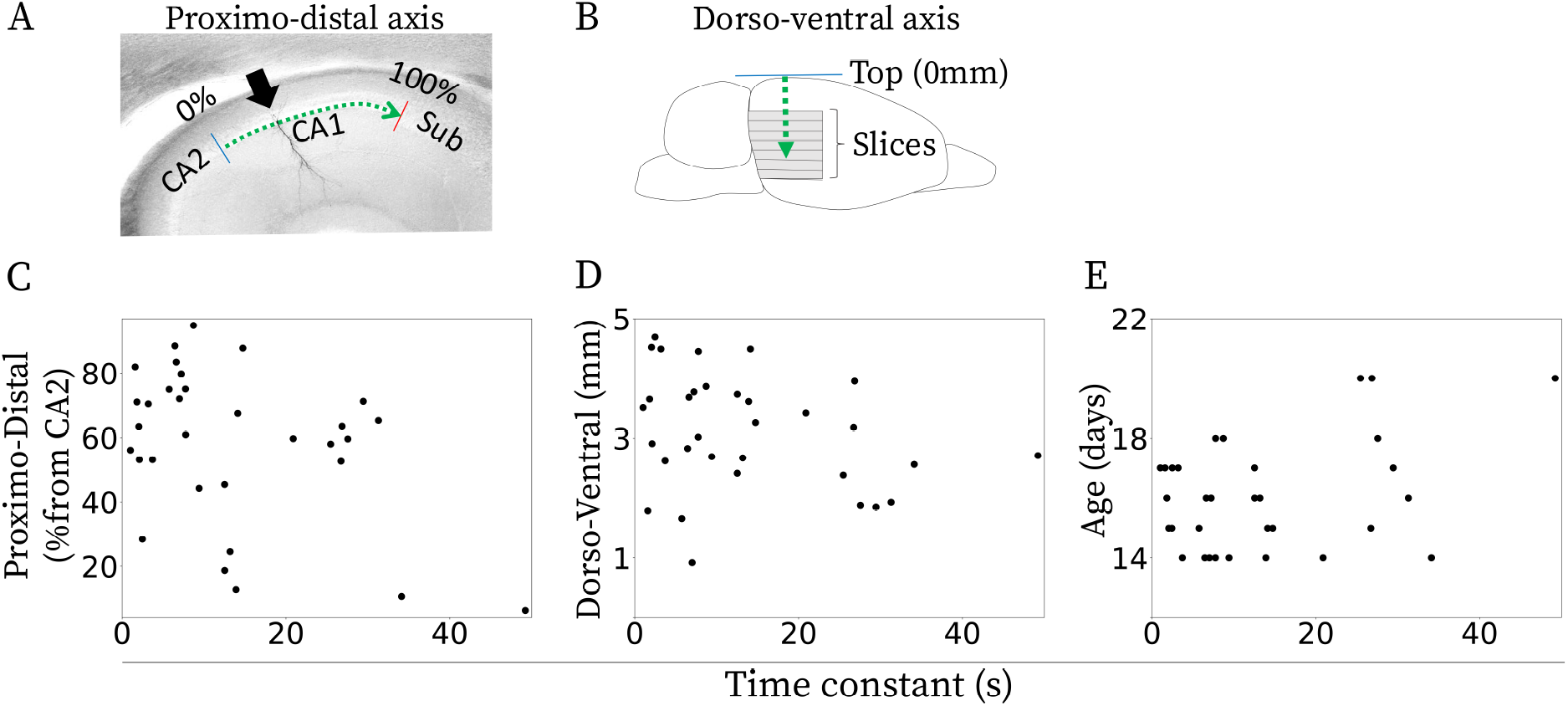
Relationship between the estimated time constant and anatomical locations and age of the neurons with exponentially decaying firing rates for all CCh concentrations combined. **A and B.** Schematic illustrating how the relative longitudinal and proximo–distal coordinates were defined in the horizontal slice preparation. (A) The blue horizontal line indicates the top of the brain reference point for dorso–ventral measurement. Slices indicate horizontal brain slices that were cut. The green arrow indicates an example dorso-ventral measurement. (B) The blue and red lines indicate CA1/CA2 and CA1/subiculum borders, respectively. The black arrow indicates the location of the recorded cell. The green arrow indicates an example proximo–distal measurement. **C.** Proximo-distal distance (% from CA2), **D.** Dorsal-ventral distance, and **E.** Age of the animal. No significant correlation was observed.

Additionally, we examined whether parameters measured during the induction of persistent firing correlated with the decay time constant. Across all CCh concentrations, no significant correlations were found with baseline membrane potential (BL), defined as the mean membrane potential during the 1s before stimulus onset (r = −0.28, p = 0.12, n = 32), or with the holding current during BL (r = 0.15, p = 0.40, n = 32). We used 1s intervals to reduce moment-to-moment noise while preserving temporal resolution. Similarly, the decay constant did not correlate with the difference between BL and the final steady state membrane potential (mean membrane potential during the last 1s of the recording; r = −0.16, p = 0.37, n = 32), nor with the difference between the immediate post-stimulus potential (mean membrane potential during the 1s after stimulation) and the final steady state membrane potential (r = −0.16, p = 0.37, n = 32).

Given the small per-condition samples, these correlation estimates are imprecise and our null findings should be interpreted cautiously.

## 4 Discussion

In this study, we report and characterize single-neuron persistent firing in hippocampal CA1 neurons in rats in the presence of synaptic blockers and CCh. Our principal finding is that, following a brief depolarizing current injection, many neurons exhibit gradually decaying firing rates spanning a wide range of time constants (approximately 1 s to 50 s). Note that the upper edge of the time constants distribution was limited by the recording duration.

These time constants span behaviorally relevant scales with many short timescales and progressively fewer long timescales. This pattern is qualitatively consistent with the compressed temporal organization reported for time-cell populations across rodents, monkeys, and humans, where later moments are represented by broader fields and by fewer cells than earlier moments rather than by a uniform tiling of time (Tiganj et al., 2017, 2018; Cruzado et al., 2020; Cao et al., 2022; Schonhaut et al., 2023). It is also consistent with monkey temporal context cells in EC, which exhibit a broad spectrum of time constants also with many short timescales and progressively fewer long timescales (Tsao et al., 2018; Bright et al., 2020).

Although temporal context cells were reported in EC (Tsao et al., 2018; Bright et al., 2020), our current investigation focused on the CA1 region primarily due to its relevance to temporal processing. Nevertheless, determining whether EC neurons exhibit a similar, or potentially more robust, intrinsic capacity for multi-scale exponential decay remains a critical question for future research.

*In vivo* ramping populations include both increasing and decreasing firing profiles. Because our analysis was anchored to the offset of a depolarizing pulse, it preferentially assayed post-stimulus relaxation back toward baseline, making down-ramping responses the expected dominant motif. The delayed-peak and other non-monotonic cells excluded from the main exponential analysis may therefore reflect neighboring dynamical regimes that are more relevant to up-ramping activity in the intact brain.

The observation that neural firing was not characterized by a stable firing rate, but by gradually changing activity, indicates that even in the presence of synaptic blockers, neurons encoded temporal information. In the absence of synaptic blockers, some of these neurons would presumably receive inputs that are modulated by the identity of the input stimuli. Coupled with the temporal modulation, this would enable simultaneous coding of “what” happened “when” in the working memory of the recent past. Neurons selective to stimulus identity and elapsed time have been reported in monkey prefrontal cortex and hippocampus during the delayed-match-to-sample task (Tiganj et al., 2018; Cruzado et al., 2020) and in bat hippocampus during social exploration (named “contextual time cells”) (Omer et al., 2023).

We also cannot exclude that some neurons would be better parameterized by oscillatory or burst-like models. This is biologically plausible because CA1 pyramidal neurons can exhibit muscarinic conditional pacemaking and cholinergically driven rhythmic activity under appropriate conditions (García-Muñoz et al., 1993; Yamada-Hanff and Bean, 2013). Visual inspection indicates that a small number of neurons showed visible oscillatory modulation superimposed on an overall decay envelope, but we did not observe a large, clearly separable class whose dominant post-stimulus pattern was pure periodic bursting (see Supplementary Material for firing profiles of all the neurons in the dataset).

### 4.1 Multi-scale exponential decay as a basis of computational models of memory and learning

We focused our analysis on exponentially decaying firing activity because such kernels play a central role in computational models of episodic memory. In Temporal Context Model (TCM) (Howard and Kahana, 2002) and its generalized version, Context Maintenance and Retrieval (CMR) model (Polyn et al., 2009), temporal context is not a single trace that simply decays from the beginning of an episode. Rather, it is a recency-weighted state that is continuously updated by ongoing experience and later reinstated at retrieval. In this framework, individual events perturb or reset context, and recent events have a larger influence than remote ones. Extensions of these models replace a single decay rate with a spectrum of exponential kernels, allowing the system to approximate an integral transform of the recent past and to represent temporal patterns over multiple scales (Shankar and Howard, 2012; Howard et al., 2015). From this perspective, the intrinsic decays reported here are best interpreted as candidate single-neuron kernels that could be combined across neurons to construct an evolving temporal context signal.

While our *in vitro* experiments did not directly test downstream readout, theoretical frameworks provide insight into how this temporal information might be decoded *in vivo*. In a freely behaving animal, the brain must extract this information from a heterogeneous population containing varying decay profiles, including the linear and multi-phase decays we observed. Downstream networks do not require *a priori* knowledge of these distinct individual profiles. Recent computational work demonstrates that biologically plausible, self-supervised learning mechanisms can naturally discover the optimal readout weights from exactly this kind of diverse population, resulting in a set of sequentially activated time cells (Alipour et al., 2025).

A neurally plausible implementation of this readout step has been proposed using off-center/on-surround weighting across neighboring time constants (Liu et al., 2019), a common connectivity motif in cortical and hippocampal microcircuits (Pouille and Scanziani, 2001). Intuitively, subtracting the activity of neurons with gradually decreasing firing rates results in a transient, bell-shaped receptive field whose peak is proportional to the time constants of the input neurons. Within the Laplace framework, this inhibitory differencing computes the inverse Laplace transform, giving rise to logarithmically compressed temporal basis functions consistent with Weber’s law (Shankar and Howard 2012).

Critically, the ingredient for power-law forgetting in the framework from Shankar and Howard (2012) and Howard et al. (2015) is that the distribution of time constants decreases as a power-law function. While the number of cells in our analysis is insufficient to argue for a particular distribution of time constants (Fig. 2A), we have illustrated that the distribution is not uniform and that the number density decreases as a function of time constant, supporting the gradual decay of memory.

If individual neurons operate in a regime where they can be considered linear systems, a population of such neurons can encode an integral transform of the recent past. A linear system would be characterized by an impulse response: the neuronal activity would be a convolution of the input signal (current injection) with the impulse response. While we established the existence of neurons with a wide range of time constants, we were unable to test whether the exponential decay can be considered the impulse response of these neurons. Since we used a single current pulse of fixed amplitude, we were unable to test the linearity of the response. Future work could investigate this by systematically manipulating the input, as well as CCh concentration, to determine whether these neurons operate within a linear regime. If single neurons truly behave as linear filters with exponential kernels, a heterogeneous population could encode rich temporal patterns. Such a mechanism could underlie a working-memory trace of “what happened when”.

In model-free and model-based reinforcement learning, efficient recursive computation of expected future reward is enabled through the Bellman equation, which requires exponential temporal discounting (Sutton and Barto, 2018; Dayan, 1993). Models with a spectrum of time constants show that integrating the exponentially discounted values leads to human-like power-law and hyperbolic discounting (Sozou, 1998; Kurth-Nelson and Redish, 2009; Tiganj et al., 2019; Momennejad and Howard, 2018; Tano et al., 2020). Exponentially decaying neurons could thus supply a substrate for encoding temporally extended value functions and guiding multi-step planning, adapting neural circuits to tasks that demand flexible action selection over a range of temporal windows.

### 4.2 Impact of CCh concentration on neural time constants

Despite significant differences across CCh concentrations, we always observed some neurons whose activity was best fit with exponentially and linearly decaying functions (Table 1). The proportion of neurons with exponentially decaying firing decreased when increasing carbachol from 5 *µM* to 20 *μ*M. One possibility is that while muscarinic receptor activation is critical for triggering persistent firing, excessive cholinergic stimulation may shift neurons toward a depolarization block or a saturated conductance state (Knauer and Yoshida, 2019). At moderate concentrations (5–10 *μ*M), the interaction between inward current that depolarized the membrane to drive persistent firing (e.g., TRPC current) and outward currents, which counteract the depolarization (e.g., SK and M currents) might have resulted in a gradually decaying persistent firing in many cells (Klink and Alonso, 1997; Egorov et al., 2002). As CCh levels rise further, however, additional or amplified inward currents may overwhelm homeostatic mechanisms, leading to prolonged depolarization and eventual block of spike generation (Tai et al., 2011; Knauer and Yoshida, 2019). Consequently, the fine-tuned balance required for a stable, exponential decay is absent in a larger fraction of neurons at 20 *μ*M.

An alternative or complementary explanation involves competing ionic or signaling processes that become dominant only under higher cholinergic drive. The same channel populations or second-messenger systems that support persistent firing at moderate CCh levels may saturate at higher concentrations. In turn, neuronal excitability and repolarization may fail to transition through the narrow parameter regime necessary for an exponential decay profile (Fransén et al., 2006). Hence, although cholinergic modulation is critical for persistent firing, these data highlight a potential nonmonotonic relationship between the degree of muscarinic receptor activation and the fraction of neurons expressing smoothly decaying activity.

When states of high cholinergic tone increase the prevalence of depolarization blocks (e.g., at 20 *μ*M CCh), a subset of cells effectively drops out of the temporal code. If the decoder is a chain-like tapped delay line (shift register), disrupting an early node would disrupt all later latencies. However, for a decoder that relies on local connectivity, like the inverse Laplace formulation, disrupting one integrator perturbs only a limited portion of the reconstructed timeline (Shankar and Howard, 2012; Liu and Howard, 2020).

An important caveat is that CCh was delivered by continuous bath application. This manipulation was useful for testing whether intrinsic mechanisms are sufficient to generate decaying persistent firing because it established a stable cholinergic permissive state, but it does not reproduce the spatially structured and temporally phasic nature of endogenous septo-hippocampal ACh release (Sarter and Lustig, 2020). Optogenetic studies in CA1 show that synaptically released ACh can recruit inhibitory interneurons, evoke rhythmic inhibitory currents, and differentially modulate Schaffer collateral and temporoammonic/entorhinal inputs through distinct muscarinic and GABA_B_/GIRK-dependent mechanisms (Nagode et al., 2014; Bell et al., 2015; Goswamee and McQuiston, 2019; Palacios-Filardo et al., 2021). Because fast ionotropic glutamatergic and GABA_*A*_-mediated transmission were blocked here, many of these circuit consequences were intentionally minimized. We therefore interpret the present results as identifying the sufficiency of an intrinsic cellular substrate rather than a full physiological account of cholinergic signaling in intact CA1. Future studies should investigate whether optogenetic cholinergic activation can also trigger decaying persistent firing over behaviorally relevant timescales.

### 4.3 Phasic cholinergic signals as event tags for encoding

In behaving animals, cholinergic activity exhibits brief, cue-locked transients that are tightly linked to whether a stimulus is detected: prefrontal acetylcholine release rises on hit trials, and bidirectional optogenetic manipulations show that suppressing these transients impairs detection, whereas generating them can increase false alarms (Parikh et al., 2007; Gritton et al., 2016). Beyond detection, these events coincide with rapid shifts in local dynamics—elevated gamma power and theta-gamma coupling during cue epochs—consistent with a fast state-switching influence on cortical processing (Howe et al., 2017). Although the precise computational role of such phasic signals remains debated (Sarter et al., 2016), their millisecond-to-second timing and trial specificity support the view that they mark behaviorally significant moments (“event boundaries”) and open brief windows of heightened excitability and plasticity.

Extending this view to hippocampal memory circuits, coordinated cholinergic release has been observed across prefrontal cortex and hippocampus on distinct timescales associated with arousal and reward, providing a route for aligning attentional and mnemonic operations around salient events (Teles-Grilo Ruivo et al., 2017). Within the hippocampal formation, cholinergic modulation biases networks toward encoding by enhancing the impact of afferent inputs and facilitating synaptic modification (Hasselmo, 2006; Teles-Grilo Ruivo and Mellor, 2013). This distinction is further supported by recent human intracranial recordings showing that muscarinic blockade with scopolamine impairs memory when present during encoding, but does not significantly impair performance when restricted to retrieval (Gedankien et al., 2025). Taken together, these findings are consistent with the possibility that brief cue-evoked cholinergic bursts could initiate or reset the intrinsic exponential decays expressed in CA1 under muscarinic activation, effectively “stamping” discrete events with a temporal tag while coordinating cortical-hippocampal processing to bind what happened to when it happened (Parikh et al., 2007; Gritton et al., 2016; Howe et al., 2017; Teles-Grilo Ruivo et al., 2017; Hasselmo, 2006; Teles-Grilo Ruivo and Mellor, 2013; Sarter et al., 2016).

### 4.4 Integration with extrahippocampal signals and behavioral state

Although we tested these neurons in a controlled slice preparation, the wide distribution of decay time constants across cells and conditions suggests a natural substrate for flexible temporal coding in the intact brain. Indeed, recent in vivo evidence supports the involvement of intrinsic persistent firing in working memory (Saber Marouf et al., 2025). In behaving animals, external inputs from frontal or parahippocampal regions could reset or modulate a neuron’s exponential decay, thus linking intrinsic cellular timescales with ongoing behavioral states (e.g., exploration, alertness, or consummatory behavior). This interaction may be particularly important when the task involves contextual changes on multi-second or minute-long intervals; exponential decays at different rates could quickly adapt to reflect new reference points for “time since event”, allowing hippocampal circuits to anchor retrieval processes or prospective encoding. Future experiments combining selective neuromodulatory manipulations and *in vivo* recordings during memory tasks would help reveal how internal dynamics, evident even in isolated neurons, interface with network-level processes to govern the timing of hippocampal-dependent behaviors.

Furthermore, these intrinsic cellular timescales must ultimately interface with the highly dynamic neuromodulatory environment of the intact brain. Recent *in vivo* studies demonstrate that hippocampal cholinergic tone is not strictly tonic; rather, acetylcholine release positively correlates with locomotion speed (Moghadam et al., 2025; Xuan et al., 2025). Previous computational work has proposed that cholinergic signals could modulate the time constants in the gradually changing activity of individual neurons (Tiganj et al., 2015). If the time constants are effectively modulated by the animal’s speed, then exponentially decaying functions of time become exponentially decaying functions of traveled distance. Thus, the same readout mechanism that can decode time from a population of neurons with gradually changing firing rates can also decode distance if the time constants are dynamically modulated by the speed (Howard et al., 2014). Fluctuations in cholinergic tone could allow the same intrinsic cellular machinery to track both time and space, consistent with neural data showing that hippocampal neurons can flexibly track time, distance, or both (Kraus et al., 2013).

## Data availability

The data that support the findings of this study are available from the corresponding author upon reasonable request. Supplementary Material contains plots of the firing profiles of all the neurons in the dataset.

## Acknowledgments

We gratefully acknowledge support from the National Institutes of Health’s National Institute on Aging, grant 5R01AG076198-02 and Deutsche Forschungsgemeinschaft Projects YO177/4-1, YO177/4-3 and YO177/7-1.

## References

Affan, R. O., Bright, I. M., Pemberton, L. N., Cruzado, N. A., Scott, B. B., and Howard, M. W. (2025). Ramping dynamics in the frontal cortex unfold over multiple timescales during motor planning. Journal of neurophysiology.

Alipour, A., James, T. W., Brown, J. W., and Tiganj, Z. (2025). Self-supervised learning of scale-invariant neural representations of space and time. Journal of Computational Neuroscience, 53(1):131—162.

Bell, L. A., Bell, K. A., and McQuiston, A. R. (2015). Activation of muscarinic receptors by ach release in hippocampal ca1 depolarizes vip but has varying effects on parvalbumin-expressing basket cells. The Journal of physiology, 593(1):197–215.

Brahimi, Y., Knauer, B., Price, A. T., Valero-Aracama, M. J., Reboreda, A., Sauvage, M., and Yoshida, M. (2023). Persistent firing in hippocampal ca1 pyramidal cells in young and aged rats. eneuro, 10(3).

Bright, I. M., Meister, M. L., Cruzado, N. A., Tiganj, Z., Buffalo, E. A., and Howard, M. W. (2020). A temporal record of the past with a spectrum of time constants in the monkey entorhinal cortex. Proceedings of the National Academy of Sciences, 117(33):20274–20283.

Cao, R., Bladon, J. H., Charczynski, S. J., Hasselmo, M. E., and Howard, M. W. (2022). Internally generated time in the rodent hippocampus is logarithmically compressed. Elife, 11:e75353.

Cao, R., Bright, I. M., and Howard, M. W. (2024). Ramping cells in the rodent medial prefrontal cortex encode time to past and future events via real laplace transform. Proceedings of the National Academy of Sciences, 121(38):e2404169121.

Cruzado, N. A., Tiganj, Z., Brincat, S. L., Miller, E. K., and Howard, M. W. (2020). Conjunctive representation of what and when in monkey hippocampus and lateral prefrontal cortex during an associative memory task. Hippocampus, 30(12):1332–1346.

Dayan, P. (1993). Improving generalization for temporal difference learning: The successor representation. Neural computation, 5(4):613–624.

Egorov, A. V., Hamam, B. N., Fransén, E., Hasselmo, M. E., and Alonso, A. A. (2002). Graded persistent activity in entorhinal cortex neurons. Nature, 420(6912):173–178.

Eichenbaum, H. (2014). Time cells in the hippocampus: a new dimension for mapping memories. Nature Reviews Neuroscience, 15(11):732–744.

Fortin, N. J., Agster, K. L., and Eichenbaum, H. B. (2002). Critical role of the hippocampus in memory for sequences of events. Nature neuroscience, 5(5):458–462.

Fransén, E., Tahvildari, B., Egorov, A. V., Hasselmo, M. E., and Alonso, A. A. (2006). Mechanism of graded persistent cellular activity of entorhinal cortex layer v neurons. Neuron, 52(5):735–744.

García-Muñoz, A., Barrio, L. C., and Buño, W. (1993). Membrane potential oscillations in ca1 hippocampal pyramidal neurons in vitro: intrinsic rhythms and fluctuations entrained by sinusoidal injected current. Experimental brain research, 97(2):325–333.

Gedankien, T., Kriegel, J., Zabeh, E., McDonagh, D., Lega, B., and Jacobs, J. (2025). Cholinergic blockade reveals a role for human hippocampal theta in memory encoding but not retrieval. eLife, 14.

Goswamee, P. and McQuiston, A. R. (2019). Acetylcholine release inhibits distinct excitatory inputs onto hippocampal ca1 pyramidal neurons via different cellular and network mechanisms. Frontiers in Cellular Neuroscience, 13:267.

Gritton, H. J., Howe, W. M., Mallory, C. S., Hetrick, V. L., Berke, J. D., and Sarter, M. (2016). Cortical cholinergic signaling controls the detection of cues. Proceedings of the National Academy of Sciences of the USA, 113(8):E1089–E1097.

Hasselmo, M. E. (2006). The role of acetylcholine in learning and memory. Current Opinion in Neurobiology, 16(6):710–715.

Howard, M. W. and Eichenbaum, H. (2013). The hippocampus, time, and memory across scales. Journal of experimental psychology: General, 142(4):1211.

Howard, M. W. and Kahana, M. J. (2002). A distributed representation of temporal context. Journal of mathematical psychology, 46(3):269–299.

Howard, M. W., MacDonald, C. J., Tiganj, Z., Shankar, K. H., Du, Q., Hasselmo, M. E., and Eichenbaum, H. (2014). A unified mathematical framework for coding time, space, and sequences in the hippocampal region. Journal of Neuroscience, 34(13):4692–4707.

Howard, M. W., Shankar, K. H., Aue, W. R., and Criss, A. H. (2015). A distributed representation of internal time. Psychological review, 122(1):24.

Howe, W. M., Gritton, H. J., Lusk, N. A., Roberts, E. A., Hetrick, V. L., Berke, J. D., and Sarter, M. (2017). Acetylcholine release in prefrontal cortex promotes gamma oscillations and theta-gamma coupling during cue detection. Journal of Neuroscience, 37(12):3215–3230.

Hsieh, L.-T., Gruber, M. J., Jenkins, L. J., and Ranganath, C. (2014). Hippocampal activity patterns carry information about objects in temporal context. Neuron, 81(5):1165–1178.

Kim, J., Ghim, J.-W., Lee, J. H., and Jung, M. W. (2013). Neural correlates of interval timing in rodent prefrontal cortex. Journal of Neuroscience, 33(34):13834–13847.

Klink, R. and Alonso, A. (1997). Muscarinic modulation of the oscillatory and repetitive firing properties of entorhinal cortex layer ii neurons. Journal of Neurophysiology, 77(4):1813–1828.

Knauer, B., Jochems, A., Valero-Aracama, M. J., and Yoshida, M. (2013). Long-lasting intrinsic persistent firing in rat ca1 pyramidal cells: A possible mechanism for active maintenance of memory. Hippocampus, 23(9):820–831.

Knauer, B. and Yoshida, M. (2019). Switching between persistent firing and depolarization block in individual rat ca1 pyramidal neurons. Hippocampus, 29(9):817–835.

Kraus, B. J., Robinson, R. J., White, J. A., Eichenbaum, H., and Hasselmo, M. E. (2013). Hippocampal “time cells”: time versus path integration. Neuron, 78(6):1090–1101.

Kurth-Nelson, Z. and Redish, A. D. (2009). Temporal-difference reinforcement learning with distributed representations. PLoS One, 4(10):e7362.

Lehn, H., Steffenach, H.-A., van Strien, N. M., Veltman, D. J., Witter, M. P., and Håberg, A. K. (2009). A specific role of the human hippocampus in recall of temporal sequences. Journal of Neuroscience, 29(11) :3475–3484.

Liu, Y. and Howard, M. W. (2020). Generation of scale-invariant sequential activity in linear recurrent networks. Neural Computation, 32(7):1379–1407.

Liu, Y., Tiganj, Z., Hasselmo, M. E., and Howard, M. W. (2019). A neural microcircuit model for a scalable scale-invariant representation of time. Hippocampus, 29(3):260–274.

MacDonald, C. J., Lepage, K. Q., Eden, U. T., and Eichenbaum, H. (2011). Hippocampal “time cells” bridge the gap in memory for discontiguous events. Neuron, 71(4):737–749.

Masse, N. Y., Yang, G. R., Song, H. F., Wang, X.-J., and Freedman, D. J. (2019). Circuit mechanisms for the maintenance and manipulation of information in working memory. Nature neuroscience, 22(7):1159–1167.

Moghadam, F. F., Gutiérrez-Guzmán, B. E., Zheng, X., Parsa, M., Hozyen, L. M., and Dannenberg, H. (2025). Cholinergic dynamics in the septo-hippocampal system provide phasic multiplexed signals for spatial novelty and correlate with behavioral states. Journal of Neuroscience, 45(41).

Momennejad, I. and Howard, M. W. (2018). Predicting the future with multi-scale successor representations. BioRxiv, page 449470.

Morcos, A. S. and Harvey, C. D. (2016). History-dependent variability in population dynamics during evidence accumulation in cortex. Nature neuroscience, 19(12):1672–1681.

Nagode, D. A., Tang, A.-H., Yang, K., and Alger, B. E. (2014). Optogenetic identification of an intrinsic cholinergically driven inhibitory oscillator sensitive to cannabinoids and opioids in hippocampal ca1. The Journal of physiology, 592(1):103–123.

Narayanan, N. S. (2016). Ramping activity is a cortical mechanism of temporal control of action. Current opinion in behavioral sciences, 8:226–230.

Omer, D. B., Las, L., and Ulanovsky, N. (2023). Contextual and pure time coding for self and other in the hippocampus. Nature neuroscience, 26(2):285–294.

Palacios-Filardo, J., Udakis, M., Brown, G. A., Tehan, B. G., Congreve, M. S., Nathan, P. J., Brown, A. J., and Mellor, J. R. (2021). Acetylcholine prioritises direct synaptic inputs from entorhinal cortex to cal by differential modulation of feedforward inhibitory circuits. Nature communications, 12(1):5475.

Parikh, V., Kozak, R., Martinez, V., and Sarter, M. (2007). Prefrontal acetylcholine release controls cue detection on multiple timescales. Neuron, 56(1):141–154.

Pastalkova, E., Itskov, V., Amarasingham, A., and Buzsáki, G. (2008). Internally generated cell assembly sequences in the rat hippocampus. Science, 321(5894):1322–1327.

Polyn, S. M., Norman, K. A., and Kahana, M. J. (2009). A context maintenance and retrieval model of organizational processes in free recall. Psychological review, 116(1):129.

Pouille, F. and Scanziani, M. (2001). Enforcement of temporal fidelity in pyramidal cells by somatic feed-forward inhibition. Science, 293(5532):1159–1163.

Saber Marouf, B., Reboreda, A., Theissen, F., Kaushik, R., Sauvage, M., Dityatev, A., and Yoshida, M. (2025). Single neurons act as a memory buffer for space. bioRxiv.

Sarter, M. and Lustig, C. (2020). Forebrain cholinergic signaling: wired and phasic, not tonic, and causing behavior. Journal of Neuroscience, 40(4):712–719.

Sarter, M., Lustig, C., Berry, A. S., Gritton, H., Howe, W. M., and Parikh, V. (2016). What do phasic cholinergic signals do? Neurobiology of Learning and Memory, 130:135–141.

Schonhaut, D. R., Aghajan, Z. M., Kahana, M. J., and Fried, I. (2023). A neural code for time and space in the human brain. Cell Reports, 42(11).

Shankar, K. H. and Howard, M. W. (2012). A scale-invariant internal representation of time. Neural Computation, 24(1):134–193.

Sozou, P. D. (1998). On hyperbolic discounting and uncertain hazard rates. Proceedings of the Royal Society of London. Series B: Biological Sciences, 265(1409):2015–2020.

Sutton, R. S. and Barto, A. G. (2018). Reinforcement learning: An introduction. MIT press.

Tai, C., Hines, D. J., Choi, H. B., and MacVicar, B. A. (2011). Plasma membrane insertion of trpc5 channels contributes to the cholinergic plateau potential in hippocampal ca1 pyramidal neurons. Hippocampus, 21(9):958–967.

Tano, P., Dayan, P., and Pouget, A. (2020). A local temporal difference code for distributional reinforcement learning. Advances in neural information processing systems, 33:13662–13673.

Teles-Grilo Ruivoy, L. M., Baker, K. L., Conway, M. W., Kinsley, P. J., Gilmour, G., Phillips, K. G., Isaac, J. T. R., Lowry, J. P., and Mellor, J. R. (2017). Coordinated acetylcholine release in prefrontal cortex and hippocampus is associated with arousal and reward on distinct timescales. Cell Reports, 18(4):905–917.

Teles-Grilo Ruivo, L. M. and Mellor, J. R. (2013). Cholinergic modulation of hippocampal network function. Frontiers in Synaptic Neuroscience, 5:2.

Tiganj, Z., Cromer, J. A., Roy, J. E., Miller, E. K., and Howard, M. W. (2018). Compressed timeline of recent experience in monkey lateral prefrontal cortex. Journal of cognitive neuroscience, 30(7):935–950.

Tiganj, Z., Gershman, S. J., Sederberg, P. B., and Howard, M. W. (2019). Estimating scale-invariant future in continuous time. Neural Computation, 31(4):681–709.

Tiganj, Z., Hasselmo, M. E., and Howard, M. W. (2015). A simple biophysically plausible model for long time constants in single neurons. Hippocampus, 25(1):27–37.

Tiganj, Z., Jung, M. W., Kim, J., and Howard, M. W. (2017). Sequential firing codes for time in rodent medial prefrontal cortex. Cerebral Cortex, 27(12):5663–5671.

Tsao, A., Sugar, J., Lu, L., Wang, C., Knierim, J. J., Moser, M.-B., and Moser, E. I. (2018). Integrating time from experience in the lateral entorhinal cortex. Nature, 561(7721):57–62.

Umbach, G., Kantak, P., Jacobs, J., Kahana, M., Pfeiffer, B. E., Sperling, M., and Lega, B. (2020). Time cells in the human hippocampus and entorhinal cortex support episodic memory. Proceedings of the National Academy of Sciences, 117(45):28463–28474.

Voelker, A. R. and Eliasmith, C. (2018). Improving spiking dynamical networks: Accurate delays, higher-order synapses, and time cells. Neural computation, 30(3):569–609.

Xuan, F., Li, G., Li, Y., and Dombeck, D. A. (2025). Modulation of speed-dependent acetylcholine release in the hippocampus by spatial task engagement. Cell reports, 44(10).

Yamada-Hanff, J. and Bean, B. P. (2013). Persistent sodium current drives conditional pacemaking in ca1 pyramidal neurons under muscarinic stimulation. Journal of Neuroscience, 33(38):15011–15021.

Yoshida, M. and Alonso, A. (2007). Cell-type–specific modulation of intrinsic firing properties and subthreshold membrane oscillations by the m (kv7)-current in neurons of the entorhinal cortex. Journal of neurophysiology, 98(5):2779–2794.

Zylberberg, J. and Strowbridge, B. W. (2017). Mechanisms of persistent activity in cortical circuits: possible neural substrates for working memory. Annual review of neuroscience, 40(1):603–627.

