## Supplementary figures for "Intrinsic persistent firing in CA1 encodes elapsed time across behaviorally relevant scales"

### Supplementary Material

#### Neurons with firing rates best fit by an exponentially decaying model in the presence of 5 $\mu M$ CCh

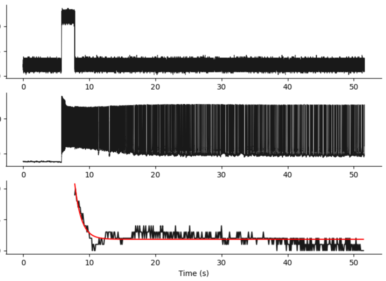


Cell ID: 1, $\tau$: 1.02

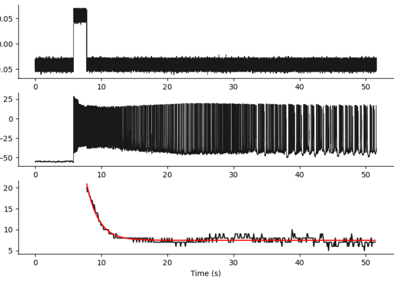


Cell ID: 2, $\tau$: 1.81

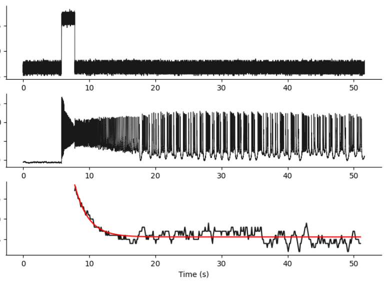


Cell ID: 3, $\tau$: 2.04

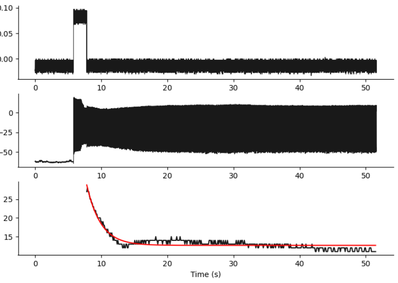


Cell ID: 4, $\tau$: 2.11

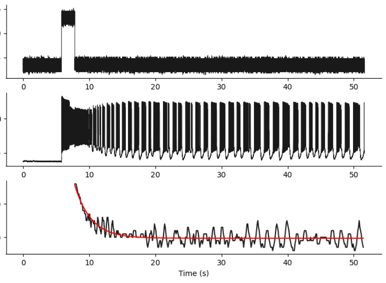


Cell ID: 5, $\tau$: 2.47

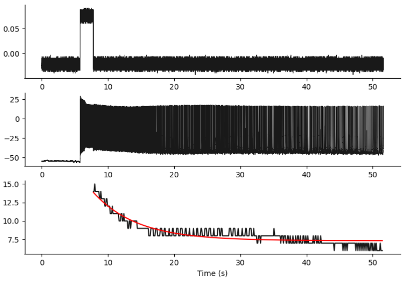


Cell ID: 6, $\tau$: 6.61

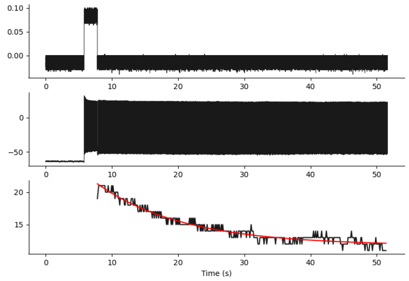


Cell ID: 7, $\tau$: 13.18

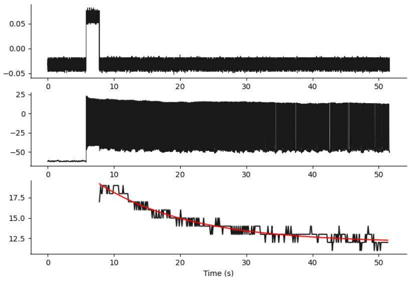


Cell ID: 8, $\tau$: 13.90

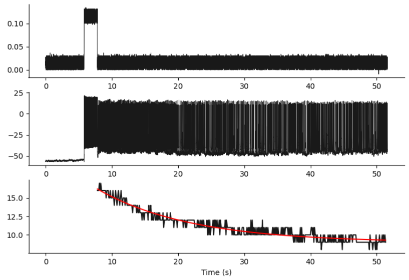


Cell ID: 9, $\tau$: 14.11

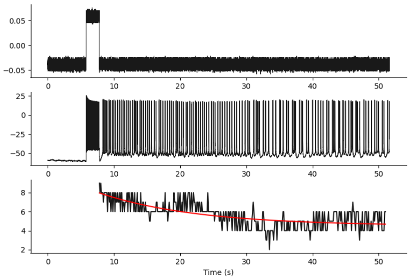


Cell ID: 10, $\tau$: 14.74

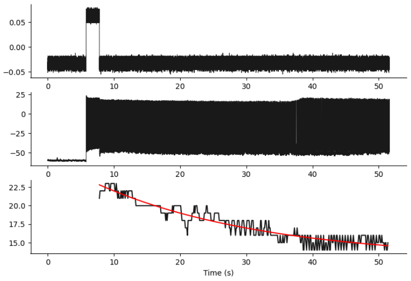


Cell ID: 11, $\tau$: 25.45

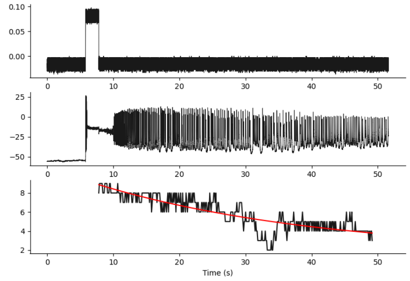


Cell ID: 12, $\tau$: 34.11

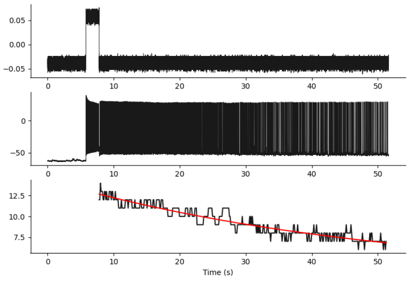


Cell ID: 13, $\tau$: 49.22

### Neurons with firing rates best fit by an exponentially decaying model in the presence of $10\ \mu M$ CCh

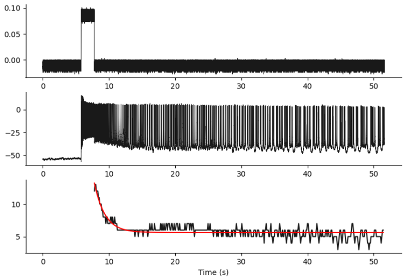


Cell ID: $14, \tau$: 1.59

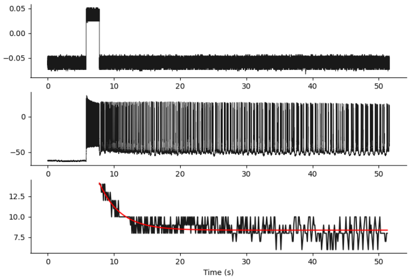


Cell ID: $15, \tau$: 3.19

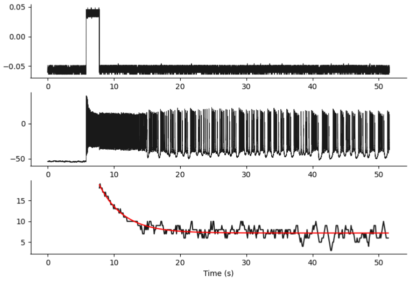


Cell ID: $16, \tau$: 3.71

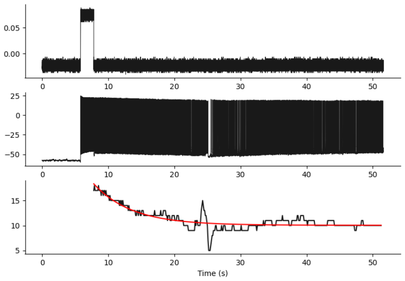


Cell ID: $17, \tau$: 5.74

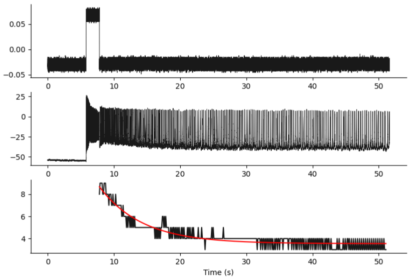


Cell ID: $18, \tau$: 6.45

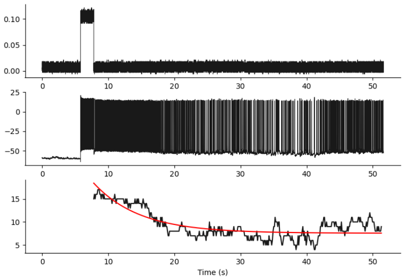


Cell ID: $19, \tau$: 7.01

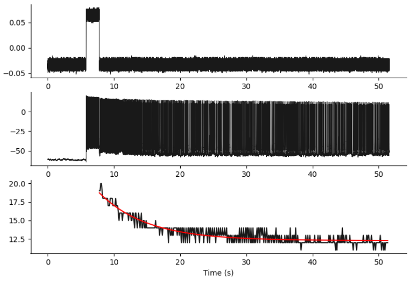


Cell ID: $20, \tau$: 7.21

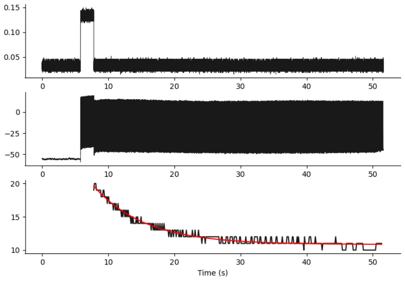


Cell ID: $21, \tau$: 7.74

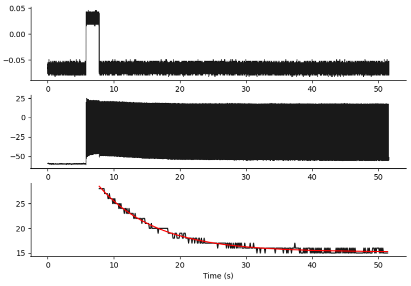


Cell ID: 22, $\tau$: 8.71

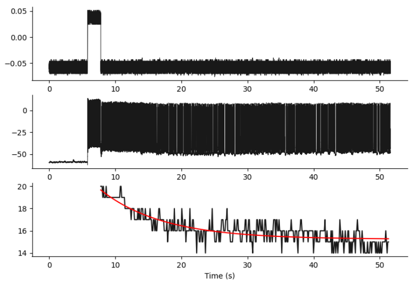


Cell ID: 23, $\tau$: 9.40

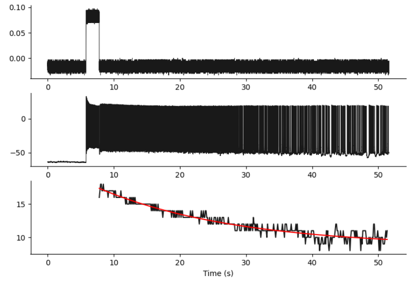


Cell ID: 24, $\tau$: 20.88

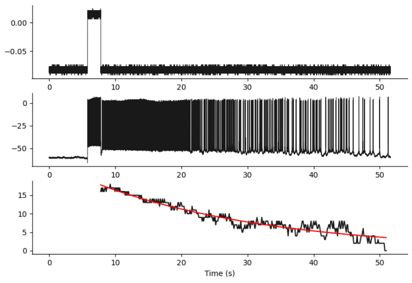


Cell ID: 25, $\tau$: 26.72

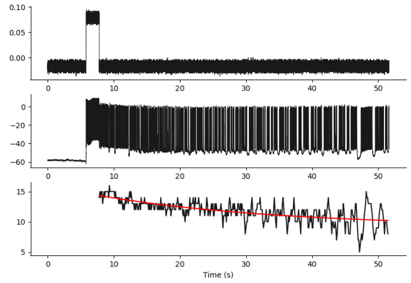


Cell ID: 26, $\tau$: 27.55

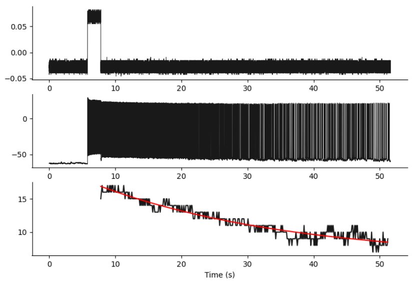


Cell ID: 27, $\tau$: 29.44

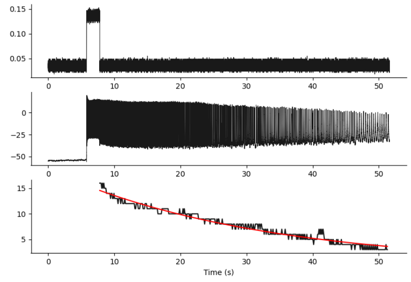


Cell ID: 28, $\tau$: 31.29

### Neurons with firing rates best fit by an exponentially decaying model in the presence of $20\ \mu M$ CCh

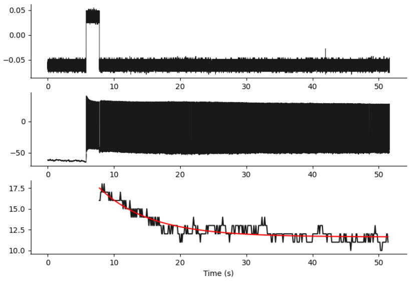


Cell ID: $29, \tau$: 7.76

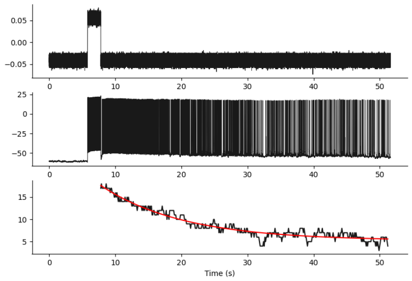


Cell ID: $30, \tau$: 12.51

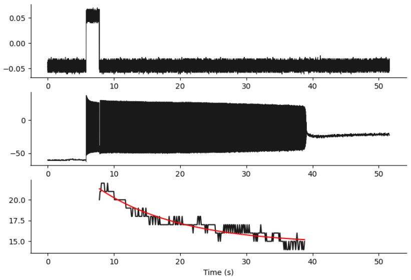


Cell ID: $31, \tau$: 12.52

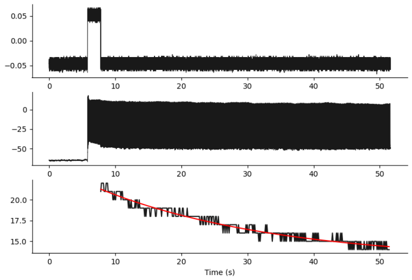


Cell ID: $32, \tau$: 26.84

#### Neurons with firing rates best fit by a linear model in the presence of 5 $\mu M$ CCh

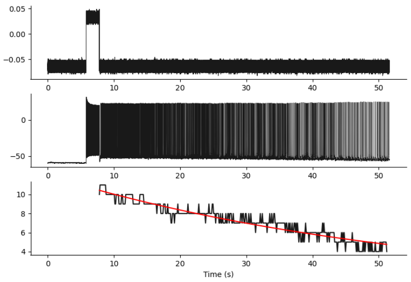

Cell ID: 33, $a$:-0.132

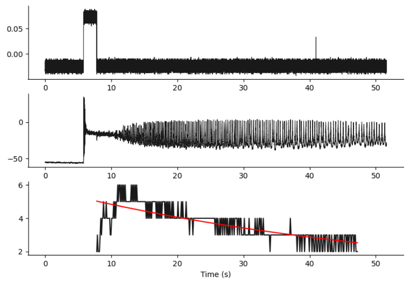

Cell ID: 34, $a$:-0.066

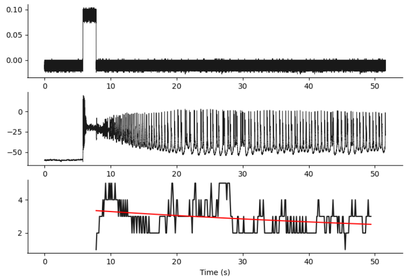

Cell ID: 35, $a$:-0.02

#### Neurons with firing rates best fit by a linear model in the presence of 10 $\mu M$ CCh

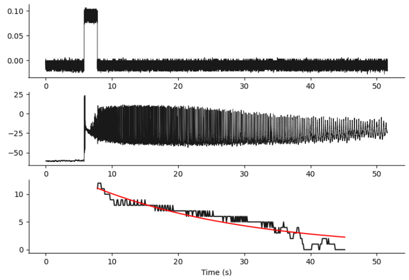


Cell ID: 36, $a$:-0.257

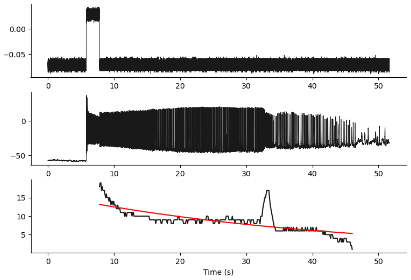


Cell ID: 37, $a$:-0.207

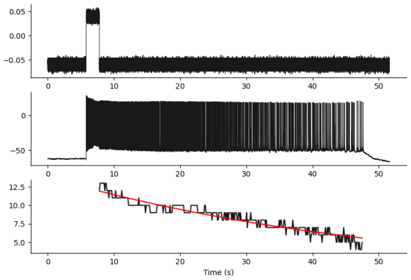


Cell ID: 38, $a$:-0.161

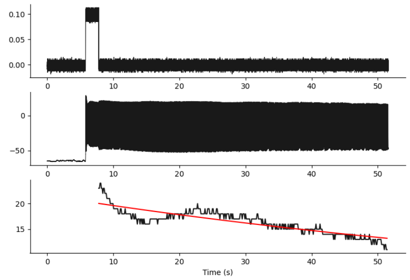


Cell ID: 39, $a$:-0.159

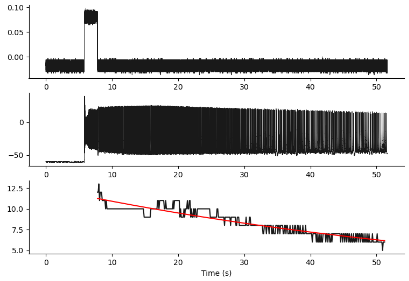


Cell ID: 40, $a$:-0.119

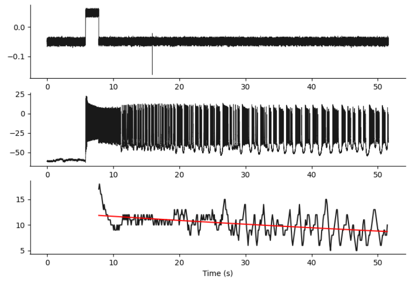


Cell ID: 41, $a$:-0.072

#### Neurons with firing rates best fit by a linear model in the presence of 20 $\mu M$ CCh

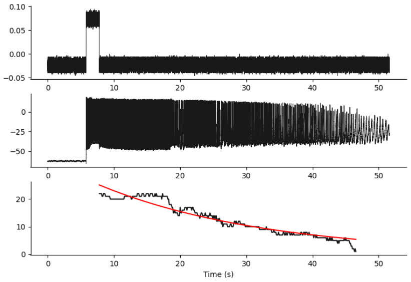


Cell ID: 42, $a$:-0.523

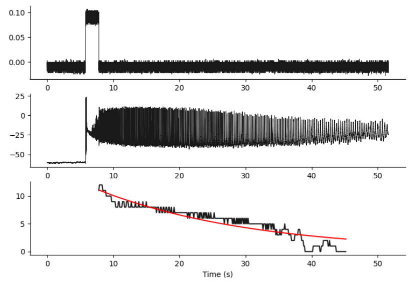


Cell ID: 43, $a$:-0.257

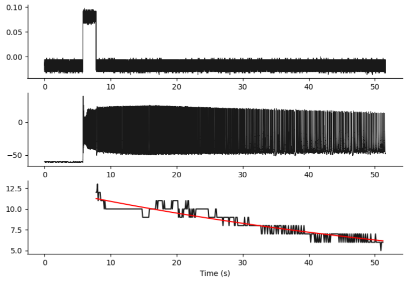


Cell ID: 44, $a$:-0.119

#### Neurons with firing rates best by a combination of linear and exponential models

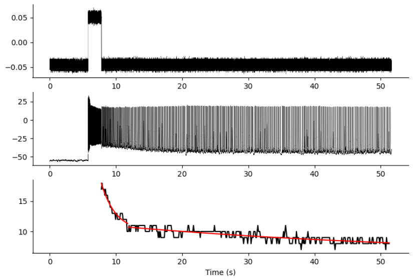


Cell ID: 45, 5 $\mu M$ CCh

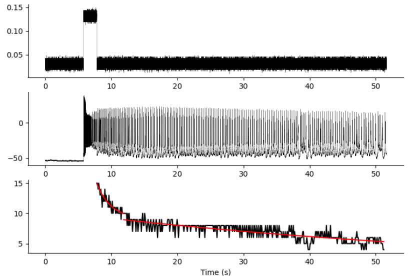


Cell ID: 46, 5 $\mu M$ CCh

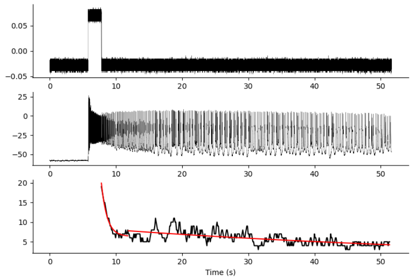


Cell ID: 47, 5 $\mu M$ CCh

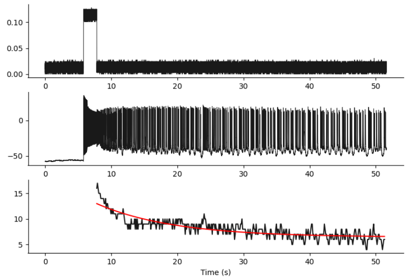


Cell ID: 48, 10 $\mu M$ CCh

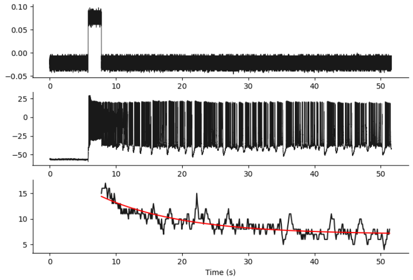


Cell ID: 49, 10 $\mu M$ CCh

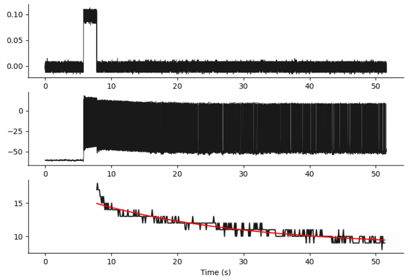


Cell ID: 50, 10 $\mu M$ CCh

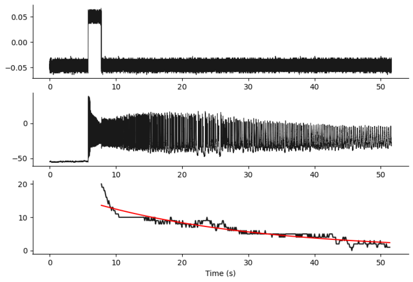


Cell ID: 51, 10 $\mu M$ CCh

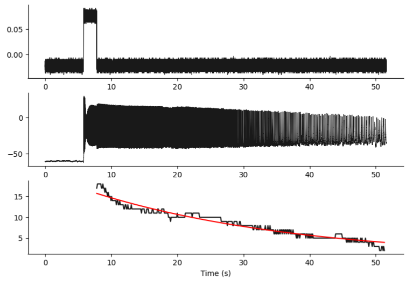


Cell ID: 52, 10 $\mu M$ CCh

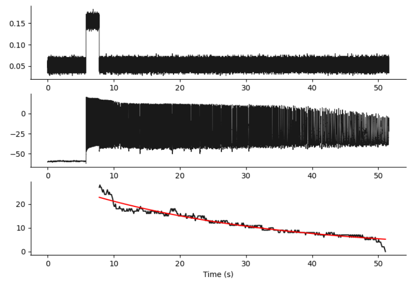


Cell ID: 53, 10 $\mu M$ CCh

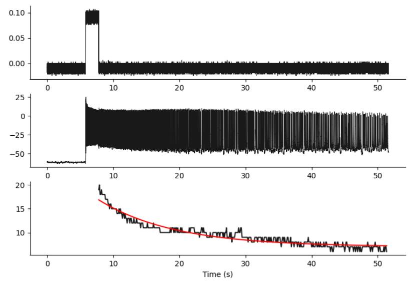


Cell ID: 54, 10 $\mu M$ CCh

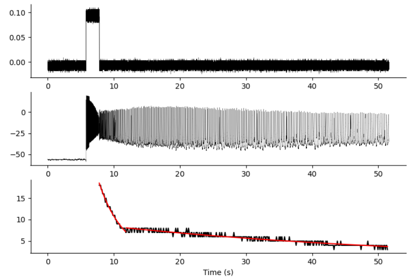


Cell ID: 55, 20 $\mu M$ CCh

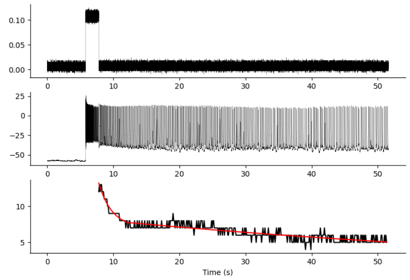


Cell ID: 56, 20 $\mu M$ CCh

### Neurons with maximum firing rate occurring after 5 s

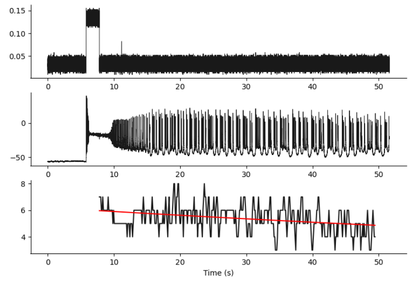


Cell ID: 57, 5 $\mu M$ CCh

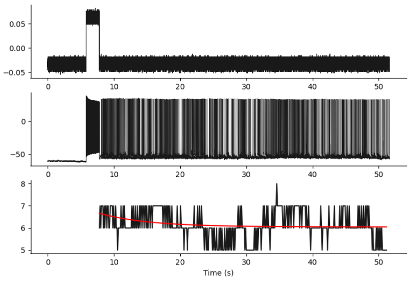


Cell ID: 58, 5 $\mu M$ CCh

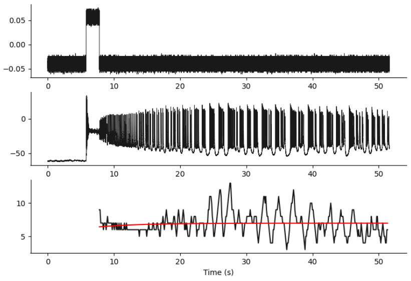


Cell ID: 59, 10 $\mu M$ CCh

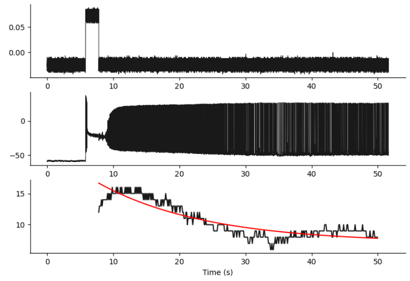


Cell ID: 60, 10 $\mu M$ CCh

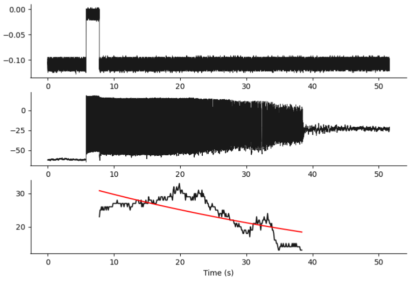


Cell ID: 61, 10 $\mu M$ CCh

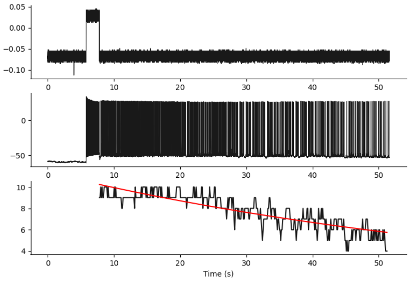


Cell ID: 62, 10 $\mu M$ CCh

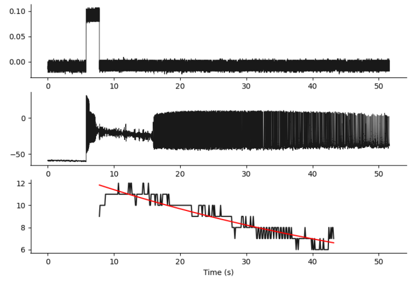


Cell ID: 63, 10 $\mu M$ CCh

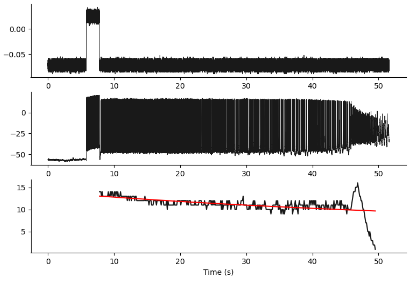


Cell ID: 64, 10 $\mu M$ CCh

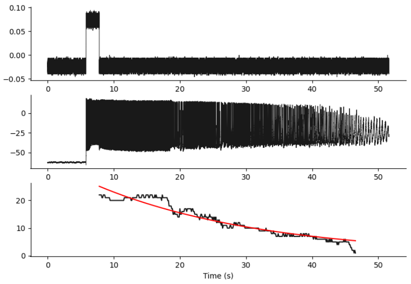


Cell ID: 65, 20 $\mu M$ CCh

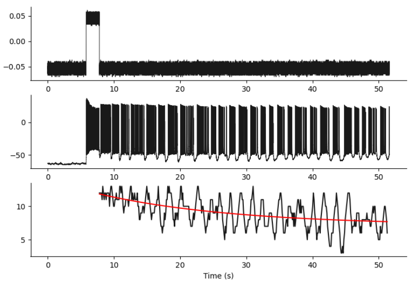


Cell ID: 66, 20 $\mu M$ CCh

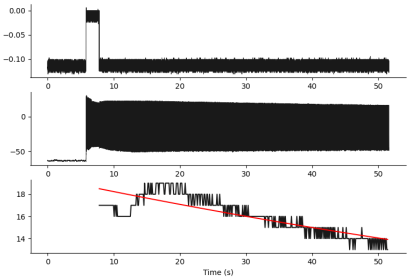


Cell ID: 67, 20 $\mu M$ CCh

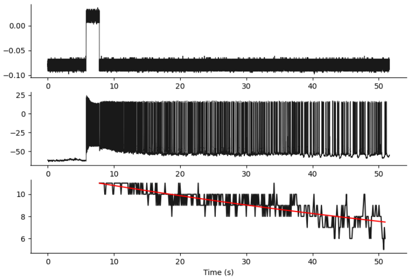


Cell ID: 68, 20 $\mu M$ CCh
